# Sex-specific hypothalamic axis disruption in the rNLS8 mouse model of amyotrophic lateral sclerosis (ALS)

**DOI:** 10.1101/2025.07.23.666305

**Authors:** Cristina Benito-Casado, Águeda Ferrer-Donato, Carmen M. Fernandez-Martos

**Affiliations:** Department of Health and Pharmaceutical Sciences, School of Pharmacy, Universidad CEU-San Pablo, CEU Universities, Madrid, Spain; Wicking Dementia Research and Education Centre, College of Health and Medicine, University of Tasmania, Hobart, Tasmania, Australia

**Keywords:** Amyotrophic lateral sclerosis (ALS), TAR DNA binding protein (TDP-43), Metabolic alterations, Leptin

## Abstract

Sex-related differences have gained increasing attention in recent years due to evidence of varying prevalence, pathophysiology, survival rates, and disease progression between males and females in amyotrophic lateral sclerosis (ALS). Differences in brain metabolism between sexes in ALS patients have also been reported; however, the specific molecular mechanisms remain poorly understood. As growing evidence supports a strong metabolic component in ALS, this study investigates alterations in leptin, one of the key regulators of metabolism, that is known to be altered during ALS progression in rNLS8 mice, a transgenic mouse model of ALS which closely recapitulates the TAR DNA-binding protein (TDP-43) pathology observed in most patients with ALS. We also examined changes in hypothalamic neuronal genes involved in metabolic regulation through food intake in rNLS8 mice. Additionally, pathological alterations in the spinal cord, a primary site of ALS pathology, were also assessed. Using molecular biology techniques we analysed the expression levels of leptin, its long receptor (Ob-Rb), and its downstream signaling pathways (Akt and STAT3) in rNLS8 mice compared to age- and sex-matched wild-type littermates. Our results revealed significant sex- and disease stage-dependent differences in leptin and Ob-Rb expression in white adipose tissue and the hypothalamus, along with increased activation of the Akt signaling pathway in the spinal cord of rNLS8 mice. These findings suggest that ALS progression differs by sex in rNLS8 mice, which may impact overall disease progression and the effectiveness of potential therapeutic interventions targeting metabolism, such as nutritional strategies that influence leptin levels.

## INTRODUCTION

Amyotrophic lateral sclerosis (ALS) is a devastating, progressive neurodegenerative disease characterized by the selective loss of motor neurons both in the brain and spinal cord [1]. ALS is considered the third most common neurodegenerative disease worldwide and is becoming a disease with a significant impact and high social awareness in many countries around the world. The majority of patients have sporadic ALS (sALS) (more than 90%) in which multiple risk factors from gene-environment interactions contribute to the disease pathogenesis [2]. In contrast, only a small subset of patients has familial ALS (fALS) (less than 10%), due to their associated genetic inheritance [2]. Over 60% of patients die within 3-5 years after diagnosis [3]. ALS affects approximately 50,000 middle-aged individuals in Europe, killing about 10,000 people. Worldwide every year 120,000 people are diagnosed with ALS. Research over the past 25 years has improved understanding of the pathophysiology of ALS, but the putative mechanisms by which mutations or gene-environment interactions cause the progressive degeneration and death of motor neurons have not been elicited yet.

Sex-related differences are increasingly recognized as a fundamental aspect of pathophysiology, prevalence, clinical presentation and disease progression in ALS [4–6]. Sex differences in ALS patients have been observed in the clinical onset of the disease, as well as in the plasma levels of adipokines such as ghrelin and leptin [7,8], which regulate metabolic pathways and are altered in metabolic disorders (e.g. obesity) and in ALS patients [9–12]. In men patients, ALS usually initiates in the lumbar tract of the spinal cord [8]. Conversely, in women patients, ALS generally begins in bulbar regions, symptoms tend to appear later, and they followed a disease progression less aggressive [8]. Moreover, lower plasma levels of the orexigenic hormone ghrelin, produced in the stomach, and the anorexigenic hormone leptin, primarily produced and secreted by the adipocytes of white adipose tissue (WAT), were found in men ALS patients [8]. This reduction in leptin levels is believed to be associated with decreased subcutaneous adiposity, a feature predominantly observed in men [13]. Indeed, the adipose tissue is a highly dynamic and heterogeneous organ, differently distributed in a sex-dependent manner [14,15], with new studies suggesting that adipocyte progenitor cells are regulated differently in female mice than in male mice [16–18]. Both in SOD1^G93A^ mice and TDP- 43^A315T^ mice models of ALS, male mice exhibit an earlier disease onset, more rapid progression and shorter lifespan compared to female mice [8,19,20]. These findings suggest that sex hormones and sex-specific metabolic regulation may play a role in modulating disease progression and possibly survival in ALS [21].

Sex differences also impact the hypothalamus, a small and key structure in the brain that regulates feeding behaviour, energy balance and metabolism. In ALS patients, hypothalamic atrophy and dysfunction have been reported [8], and emerging evidence suggests that these alterations may differ between men and women [8]. For instance, sex- specific patterns of hypothalamic gene expression and leptin signaling have been observed in animal models [22,23], potentially contributing to differences in metabolic regulation and disease progression. Agouti-related peptide (AgRP) and proopiomelanocortin (POMC) neurons, located in the arcuate nucleus in the hypothalamus, have an important role in modulating feeding behaviour [24,25]. Neuropeptide Y (NPY)/AgRP neurons are inhibited by leptin, that at the same time activate POMC/cocaine-and amphetamine-regulated transcript (CART) neurons, maintaining energy homeostasis, as it leads to satiety, and controlling body mass index (BMI) [26]. Leptin also activates the sympathetic nervous system, promoting lipolysis in WAT and thermogenesis in brown adipose tissue [27,28]. Leptin receptors are included in class I cytokine receptors, and to date six leptin receptor are known that arise from different splicing (Ob-Ra-f) [29]. Although these receptors are expressed in the hypothalamus, as well as in other extra-hypothalamic regions, leptin signals via the long isoform of the receptors (Ob-Rb) [29]. A state of leptin resistance can occur when mTOR activity is increased in POMC neurons, as it is observed in obese mice [30,31]. However, despite these insights, there remains a lack of comprehensive understanding how these sex- specific metabolic and neuroanatomical differences influence disease progression, which is crucial for developing targeted therapeutic strategies.

In this context, we examined the molecular mechanisms underlying these metabolic differences, with a particular focus on leptin signaling, hypothalamic regulation, and adipose tissue innervation in both male and female rNLS8 mice, compared to sex- and age-matched wild-type (WT) littermates, providing, to our knowledge, the first insights into significant sex-dependent differences in the regulation of these pathways during disease progression in rNLS8 mice.

## MATERIALS AND METHODS

### Animals

Forty hTDP-43ΔNLS animals were used in this study (referred to as regulatable NLS, rNLS8), both male and female and age-matched WT littermates. rNLS8 mice were produced crossing hemizygous *tetO-*hTDP-43- ΔNLS (RRID:IMSRJAX:014650) with hemizygous *NEFH*-tTA (RRID:IMSRJAX:025397) on a mixed B6/C3H background [32], at Hospital Nacional de Parapléjicos. Both rNLS8 and WT mice were grouped house under identical conditions in a 12-hour standard light/dark cycle with *ad libitum* access to water and food. Mice were fed with a standard chow containing 200 mg/kg doxycycline (dox) and switched to standard chow lacking dox at 4 weeks of age. Diets were provided by Tecklad Global diets, Evigo (IT). Mice were randomly assigned to one- or four-weeks post-dox (referred to as 1w and 4w off-dox). 1w off-dox at the onset of pathological induction of TDP-43 and 4w off-dox since the animals have been reported to present decrease muscle innervation, cortical neuron degeneration and astrogliosis [33].

hTDP-43ΔNLS expression was confirmed via PCR according to the distributor’s protocol. The maintenance and use of mice and all experimental procedures were approved by the Animal Ethics Committee of the of the National Hospital for Paraplegics (HNP) (Approval No 26/OH 2018) in accordance with the Spanish Guidelines for the Care and Use of Animals for Scientific Purposes.

### Perfusion and tissue collection

Mice were terminally injected intraperitoneally with sodium pentobarbitone (140 mg/kg) and transcardially perfused with room temperature (RT) 0.01 M phosphate buffered saline (PBS; pH 7.4). Hypothalamus, lumbar spinal cord and WAT were dissected and processed for real-time polymerase chain reaction (RT-qPCR) and western blot (WB) analysis. Samples were immediately frozen on dry ice and stores at -80 °C for later analysis.

### RNA Isolation and RT-qPCR

Total RNA was isolated from WAT and hypothalamus using the RNeasy Mini Kit (Qiagen), according to the manufacturer’s instructions. Complementary DNA (cDNA) was synthesized as described previously [34] from 1 µg and 0.5 µg of total RNA of WAT and hypothalamus, respectively. Relative quantification of *ObRb, Leptin, C/EBP*β*, PPAR*γ*, AgRP, POMC* and *NPY* was performed as described previously [35]. To amplify the cDNA of TDP-43 the following primers (Invitrogen) were used (forward *5*′*- GGGCGATGGTGTGACTGTAA-3*′ and reverse *5*′*-GCTCGTCTGGGCTTTGCTTA-3*′.

GenBank accession number: NM_145556.4). The 18S rRNA was used as a control to normalize gene expression [36]. The reactions were run on an CFX96 Real-Time System instrument and software (CFX Manager 3.0) (BioRad) according to the manufacturer’s protocol. Relative quantification for each gene was performed by the ΔΔCt method [37].

### Protein extraction and WB analysis

RIPA buffer (Sigma Aldrich) containing a cocktail of protease inhibitors (Roche) as described previously [11] was used to extract proteins from spinal cord. 20 µg of denatured protein samples from each group were electrophoresed into Bolt® Bis–Tris Plus gels (Invitrogen), transferred to PVDF membranes (BioRad) and incubated with primary antibodies [rabbit anti-ObRb (1:500; Abcam), rabbit anti-leptin (1:400, Invitro), mouse anti- STAT3 (1:500; Santa Cruz), rabbit anti-pSTAT3 (1:1000; Cell Signalling), mouse anti-Akt (1:500; Santa Cruz), rabbit anti-pAkt (1:500; Cell Signaling)] overnight. A corresponding anti-rabbit or anti-mouse horseradish peroxidase (HRP)-conjugated secondary antibody (Vector Laboratories) at dilution of 1:5000 was used as described previously [11]. Mouse anti-GAPDH (1:5000, Milipore) was used as a loading control and band intensity was measured as the integrated intensity using ImageJ software (v1.4; NIH). All data were normalized to control values on each membrane.

### Statistical Analysis

All data are presented as the mean ± standard error of the mean (SEM). Normality of datasets was assessed by Shapiro-Wilk test. Outliers were removed wit ROUT method with Q=1%. Differences between means were assessed by two-way ANOVA followed by Dunnett’s post hoc test and Tukey’s post hoc test. To compared within the same group, T- test and Mann-Whitney test were used depending on whether they had a normal distribution or not. For all statistical tests, a *p*-value of <0.05 (CI 95%) was assumed to be significant. Statistical analysis was performed using GraphPad Prism software (version 10.2.0).

## RESULTS

### Leptin levels are altered in WAT of rNLS8 mice

Given the limited insight into the status of leptin expression in the WAT of rNLS8 mice, we aimed to address this gap by performing a comprehensive molecular characterization of *leptin* and *Ob-Rb* mRNA levels in WAT during the disease progression (1w vs. 4w off-dox) in rNLS8 mice compared to sex and age-matched WT littermates, using RT-qPCR analysis. RT-qPCR analysis demonstrated marked differences in the expression profile of *leptin* and *Ob-Rb* transcripts during the disease progression in the WAT of rNLS8 mice compared to WT controls (Figure 1). As preliminary observations suggest that the clinical course of the disease may differ between male and female rNLS8 mice [38], highlighting the need for further investigation into sex-specific mechanisms in this ALS model, we have separated them for the analysis. In male rNLS8 mice, we found a significant effect of genotype (*p*=0.022) on *Ob-Rb* mRNA levels (Figure 1A). There was an up-regulation of *Ob-Rb* mRNA levels in male rNLS8 mice at 4w off-dox compared to age-matched WT littermates, and in male rNLS8 mice at 1w off-dox compared to WT controls at 1w off-dox, although it did not reach significance (Figure 1A). We found a significant effect of disease progression (*p*=0.002) on *leptin* mRNA levels (Figure 1C) in male rNLS8 mice. RT-qPCR analysis showed a significant down-regulation of *leptin* mRNA levels in male rNLS8 mice at 4w off-dox compared to WT littermates at 1w off-dox and compared to male rNLS8 mice at 1w off-dox and an up-regulation in male rNLS8 mice at 1w off-dox compared to WT controls at 4w off-dox (Figure 1C, *p*=0.013, *p*=0.047 and *p*=0.036, respectively). Conversely, in female rNLS8 mice there was no significant effect either of disease progression or genotype on *Ob-Rb* mRNA levels (Figure 1B), although there was a significant effect on disease progression (*p*=0.037) on *leptin* mRNA expression levels (Figure 1D).

**Figure 1.**
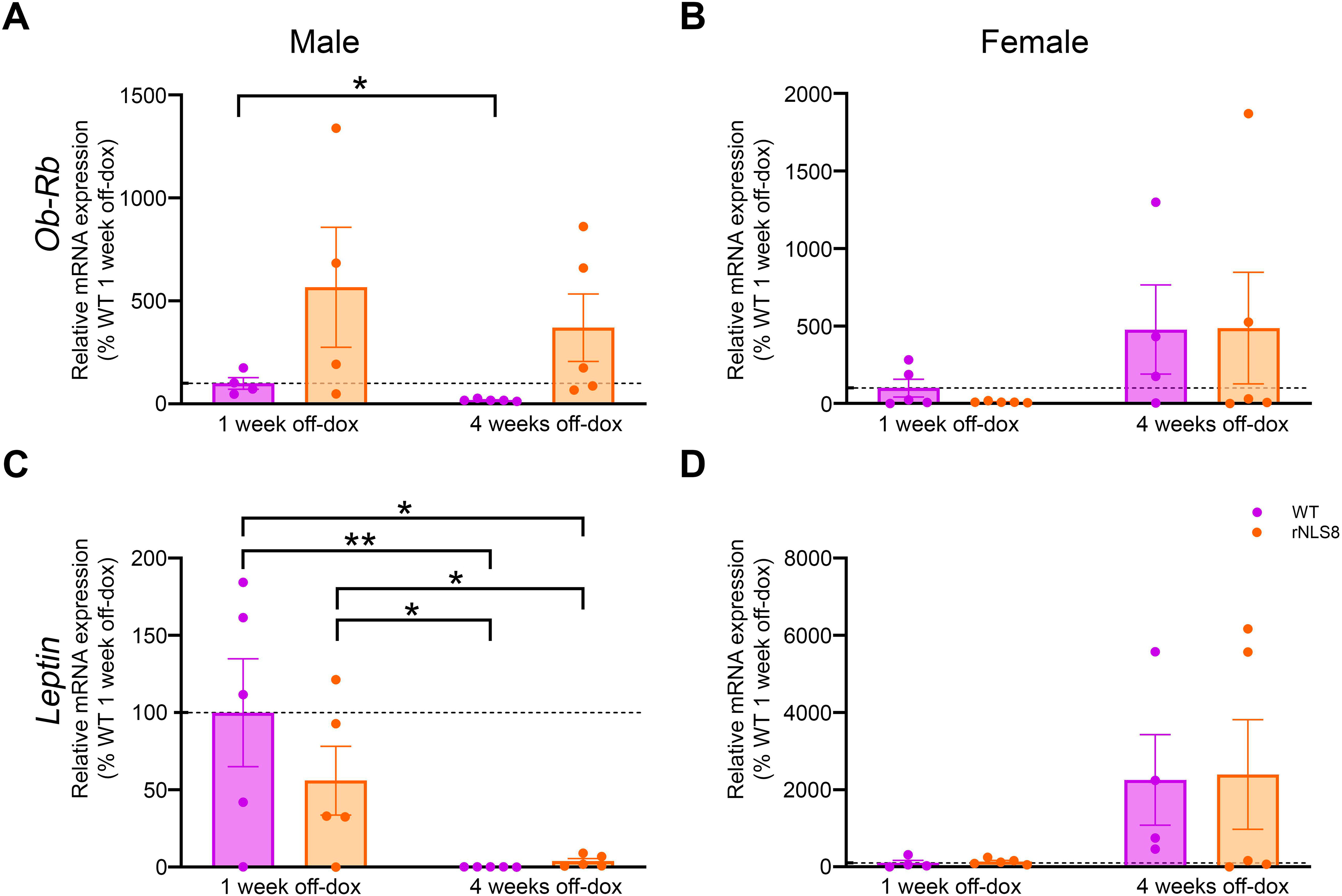
Alterations in *Ob-Rb* and *leptin* in WAT of rNLS8 mice. (A) Male *Ob-Rb*, (B) female *Ob-Rb*, (C) male *leptin* and (D) female *leptin* mRNA expression. Transcripts were assessed by RT-qPCR in rNLS8 mice compared to age- and sex-matched WT littermates at 1 and 4w off-dox. Values are expressed as the mean ± SEM for the different groups. *p<0.05, **p<0.01, ***p<0.001. Abbreviations: Abbreviations: *Ob-Rb*, long form of the leptin receptor; WAT, white adipose tissue; RT-qPCR, Reverse Transcription quantitative Polymerase Chain Reaction; WT, wild-type; 1w, 1 week; 4w, 4 weeks; dox, doxycycline.

Previous results from our group, not yet published [39], showed lower circulating leptin levels in plasma in male rNLS8 mice compared to WT controls at 4w off-dox. New results from our group confirm a significant impairment of WAT during the progression of disease in male TDP-43^A315T^ mice compared to WT controls [40]. Our preliminary unpublished data [40] indicated how these histopathological alterations are less severe in females TDP- 43^A315T^ mice. Therefore, to further elucidate the molecular alterations occurring in WAT during the progression of disease in rNLS8 mice, we analyse genes controlling adipocyte differentiation. In male rNLS8 mice, RT-qPCR analysis showed significant effect of disease progression and genotype on *C/EBP*β (Figure 2A, *p*=0.014 and *p*=0.042, respectively) and *TDP-43* mRNA expression levels (Figure 2E, *p*=0.030 and *p*=0.043, respectively), while no significant effect on neither of them was observed on *PPAR*γ mRNA (Figure 2C). *C/EBP*β and *TDP-43* expression mRNA levels were upregulated in male rNL8 mice at 1w off-dox compared to WT controls at 4w off-dox (Figure 2A, *p*=0.013 and Figure 2E, *p*=0.024, respectively). Furthermore, in female rNLS8 mice, RT-qPCR analysis showed significant effect of disease progression (*p=*0.006) and genotype (*p*=0.004) on *C/EBP*β (Figure 2B), while it only showed a significant effect of disease progression (*p*=0.021) on *PPAR*γ mRNA levels (Figure 2D). In addition, *C/EBP*β was significantly downregulated in female rNLS8 mice at 1 and 4w off-dox compared to WT controls at 4w off-dox (Figure 2B, *p*=0.002 and *p*=0.015, respectively). Finally, *PPAR*γ was significantly downregulated in female rNLS8 mice at 1w off-dox compared to WT controls at 4w off-dox (Figure 2D, *p*=0.009). No significant effect neither of disease progression nor genotype was found on *TDP-43* mRNA levels (Figure 2F), however, levels were downregulated in female rNLS8 mice at 1w off-dox compared to WT controls at 4w off-dox (Figure 2F, *p*=0.016).

**Figure 2.**
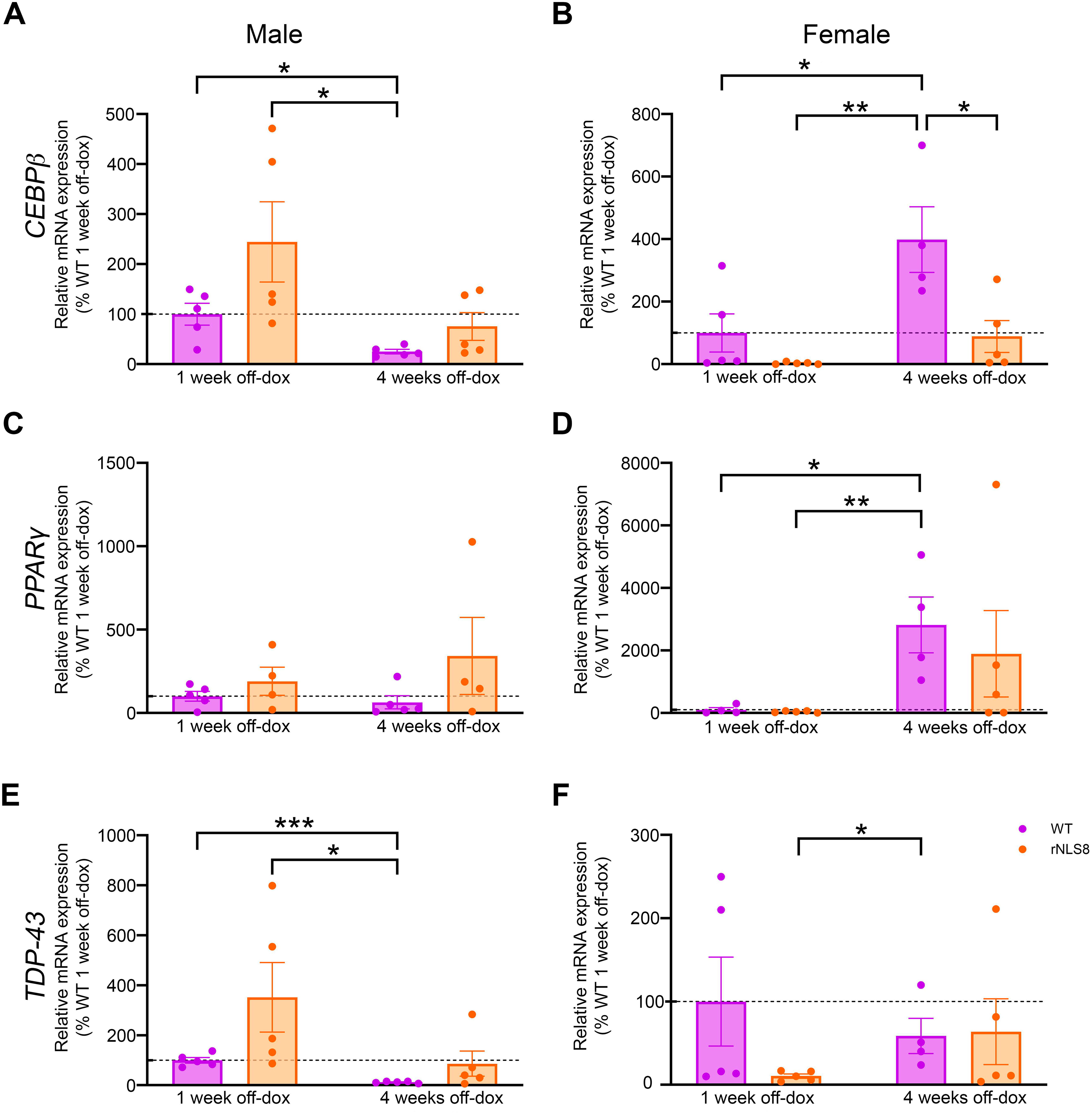
**Alterations in *C/EBP***β**, *PPAR***γ **and *TDP-43* in the WAT of rNLS8 mice.** (A) Male *C/EBP*β, (B) female *C/EBP*β, (C) male *PPAR*γ, (D) female *PPAR*γ, (E) male *TDP-43* and (F) female *TDP-43* mRNA expression. Transcripts were assessed by RT-qPCR in rNLS8 mice compared to age- and sex-matched WT littermates at 1 and 4w off-dox. Values are expressed as the mean ± SEM for the different groups. *p<0.05, **p<0.01, ***p<0.001. Abbreviations: *C/EBP*β, CCAAT/enhancer-binding protein beta; *PPAR*γ, Peroxisome proliferator-activated receptor gamma; *TDP-43*, TAR DNA-binding protein 43; WAT, white adipose tissue; RT-qPCR, Reverse Transcription quantitative Polymerase Chain Reaction; WT, wild-type; 1w, 1 week; 4w, 4 weeks; dox, doxycycline.

### Hypothalamic leptin signaling in rNLS8 mice

As the central hypothalamic leptin signaling has a critical role in promoting energy homeostasis [41], we studied the status of leptin signaling and neuropeptides in the hypothalamus of rNLS8 mice at both time points of the disease. In male rNLS8 mice, RT- qPCR analysis showed no significant effect neither of disease progression or genotype on *Ob-Rb* (Figure 3A) nor *NPY* (Figure 3C) mRNA levels, but it showed a significant effect of disease progression (*p*=0.037) on *AgRP* (Figure 3E) and on genotype (*p*=0.015) on *POMC* mRNA levels (Figure 3G). *Ob-Rb* and *POMC* mRNA levels were downregulated in the hypothalamus of male rNLS8 mice at 4w off-dox compared to WT controls at 1w off-dox (Figure 3A, *p*=0.024 and Figure 3G, *p*=0.036, respectively).

**Figure 3.**
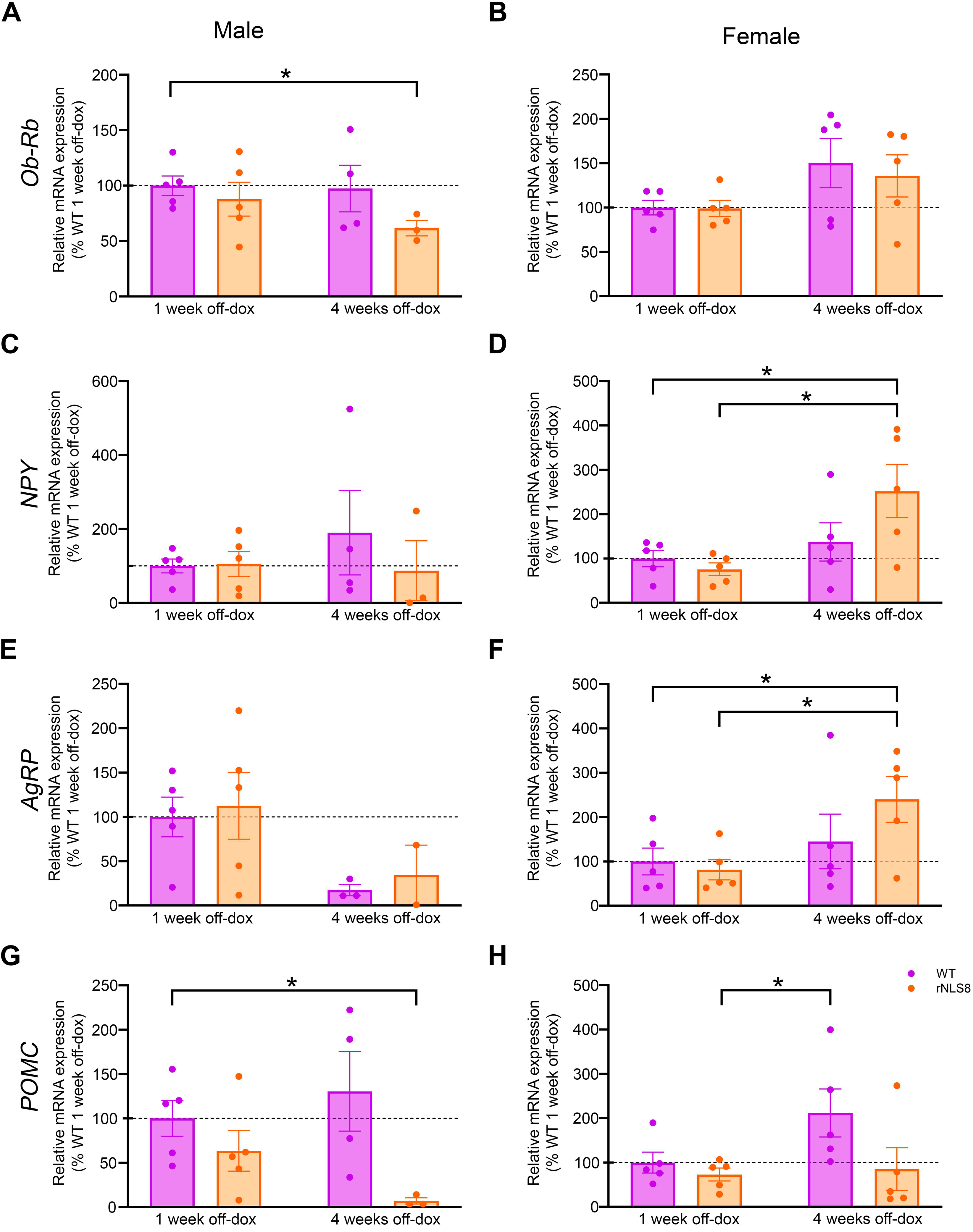
Alterations in *Ob-Rb*, *AgRP, NPY* and *POMC* in the hypothalamus of rNLS8 mice. (A) Male *Ob-Rb*, (B) female *Ob-Rb*, (C) male *AgRP*, (D) female *AgRP*, (E) male *NPY*, (F) female *NPY*, (G) male *POMC* and (H) female *POMC* mRNA expression. Transcripts were assessed by RT-qPCR in rNLS8 mice compared to age- and sex- matched WT littermates at 1 and 4w off-dox. Values are expressed as the mean ± SEM for the different groups. *p<0.05, **p<0.01, ***p<0.001. Abbreviations: *Ob-Rb*, long form of the leptin receptor; *AgRP*, Agouti-related protein; *NPY*, Neuropeptide Y; *POMC*, proopiomelanocortin; RT-qPCR, Reverse Transcription quantitative Polymerase Chain Reaction; WT, wild-type; 1w, 1 week; 4w, 4 weeks; dox, doxycycline.

In female rNLS8 mice, RT-qPCR analysis showed significant effect of disease progression on *Ob-Rb* (Figure 3B, *p*=0.038), *NPY* (Figure 3D, *p*=0.014) and *AgRP* (Figure 3F, *p*=0.035), while no significant effect neither disease progression nor genotype were showed on *POMC* mRNA levels (Figure 3H). Regarding orexigenic neuropeptides*, NPY* and *AgRP* mRNA levels were upregulated in female rNLS8 mice at 4w off-dox compared to female rNLS8 mice at 1w off-dox (Figure 3D, *p*=0.021 and Figure 3F, *p*=0.022, respectively). Finally, *POMC* mRNA levels were significant downregulated in female rNLS8 mice at 1w off-dox compared to WT controls at 4w off-dox (Figure 3H, *p*=0.039).

### Leptin signaling in the spinal cord of rNLS8 mice

As previous published results of our group demonstrated significant alterations in the protein expression levels of Ob-R and and its downstream signaling pathways (Akt and STAT3) in TDP-43^A315T^ mice [11], we investigated the protein levels of Ob-Rb and leptin, as well as the status of STAT3 (pTyr^705^-STAT3) and Akt (pSer^473^-Akt) pathways in rNLS8 mice, which is currently unexplored. WB analysis demonstrated there was a significant effect of disease progression (*p*<0.001) and genotype (*p*<0.001) on the expression profile of the Ob-Rb receptor in the spinal cord of male rNLS8 mice compared to WT controls (Figure 4A). Protein levels of the Ob-Rb receptor were significantly lower in male rNLS8 mice at 1w off-dox compared to WT control at 4w off-dox and to male rNLS8 mice at 4w off-dox (Figure 4A, *p*<0.001 and *p*=0.007, respectively) and in male rNLS8 mice at 4w off- dox compared to age-matched WT controls and to WT controls at 1w off-dox (Figure 4A, *p*<0.001 and *p*=0.022, respectively). We also found a significant effect of disease progression (*p*<0.001) and genotype (*p*=0.023) on leptin levels (Figure 4C). Leptin protein levels were lower in male rNLS8 mice at 1w off-dox compared to WT controls at 4w off-dox and to male rNLS8 mice at 4w off-dox (Figure 4C, *p*<0.001 and *p*=0.046, respectively) and in male rNLS8 at 4w off-dox compared to age-matched WT controls and to WT controls at 1w off-dox (Figure 4C, *p*=0.004 and *p*=0.019, respectively). In addition, we also found a significant effect of disease progression (*p*<0.001) on the phosphorylation levels of STAT3 protein (Figure 5A). STAT3 phosphorylation levels were significantly increased in male rNLS8 mice at 1w off-dox compared to WT controls at 4w off-dox and to male rNLS8 mice at 4w off-dox (Figure 5A, *p*<0.001 and *p*<0.001, respectively) and in male rNLS8 mice at 4w off-dox compared to WT controls at 1w off-dox (Figure 5A, *p*<0.001). Interestingly, there was a significant effect of disease progression (*p*<0.001) and genotype (*p*=0.034) on the phosphorylation levels of Akt protein (Figure 5C). At 1w off-dox, Akt phosphorylation levels of male rNLS8 mice were significantly increased compared to age-matched WT controls and to WT controls and male rNLS8 mice at 4w off-dox (Figure 5C, *p*=0.002, *p*=0.003 and *p*<0.001, respectively) and in male rNLS8 mice at 4w off-dox compared to age-matched WT controls and to WT controls at 1w off-dox (Figure 5C, *p*=0.020 and *p*<0.001, respectively).

**Figure 4.**
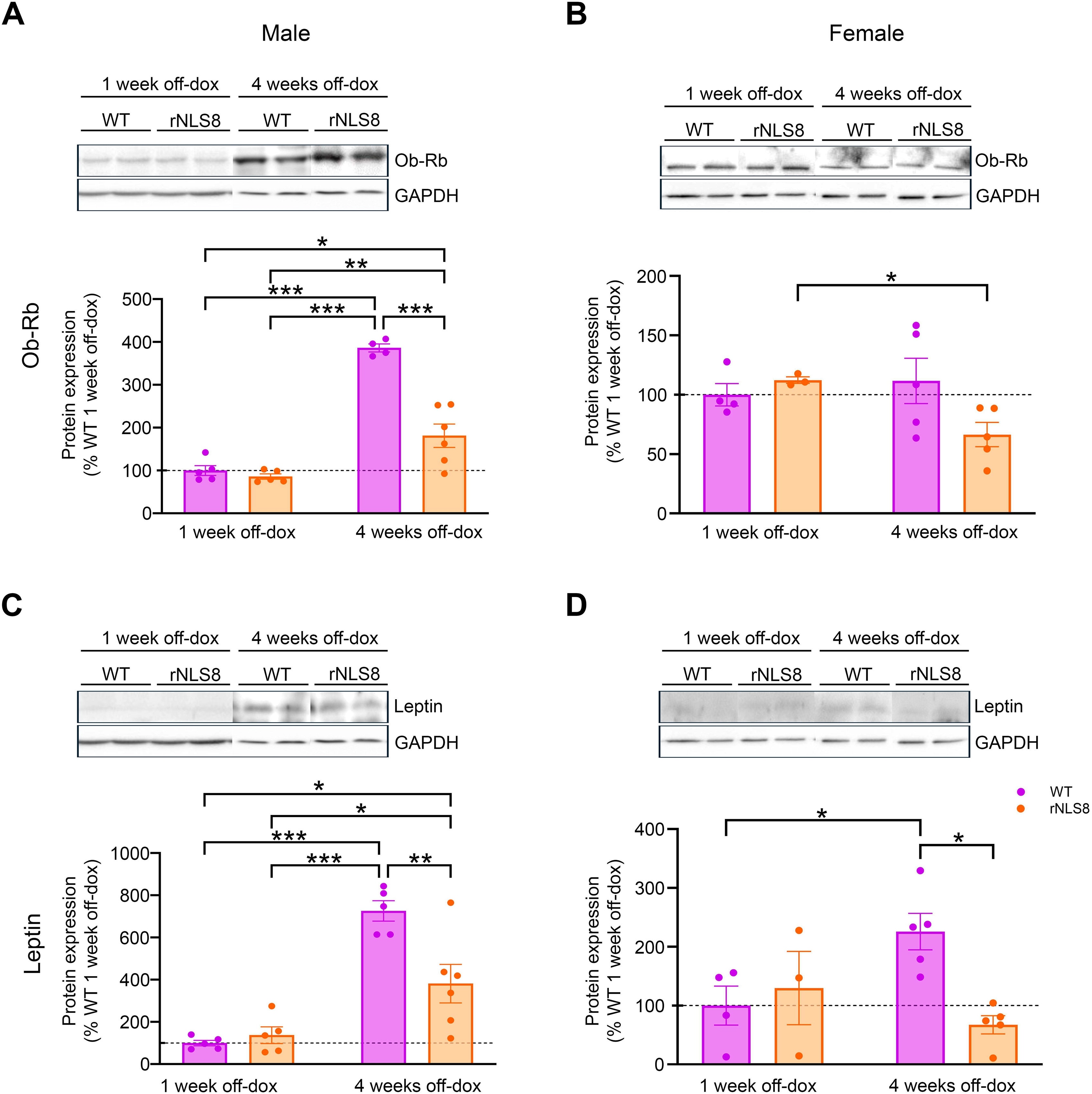
Alterations in Ob-Rb and leptin in the spinal cord of rNLS8 mice. (A) Male Ob-Rb, (B) female Ob-Rb, (C) male leptin, (D) female leptin in the spinal cord extracts of rNLS8 mice compared to age- and sex-matched WT littermates at 1 and 4w off-dox. *p<0.05, **p<0.01, ***p<0.001. Abbreviations: Ob-Rb, long form of the leptin receptor; WT, wild-type; 1w, 1 week; 4w, 4 weeks; dox, doxycycline.

**Figure 5.**
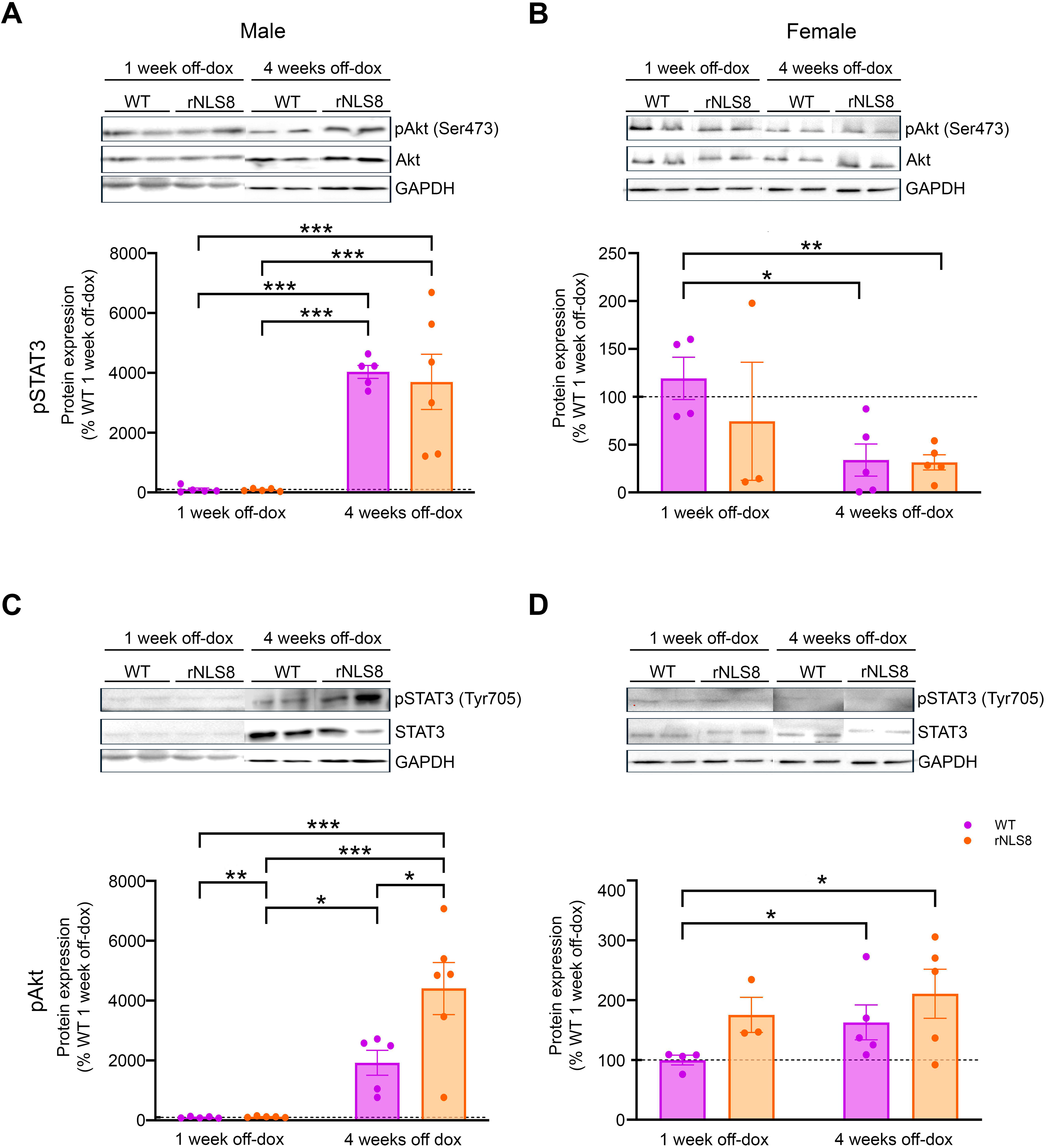
Alterations in pSTAT3 and pAkt in the spinal cord of rNLS8 mice. (A) Male pSTAT3, (B) female pSTAT3, (C) male pAkt, (D) female pAkt in the spinal cord extracts of rNLS8 mice compared to age- and sex-matched WT littermates at 1 and 4w off-dox. *p<0.05, **p<0.01, ***p<0.001. Abbreviations: pSTAT3, pTyr^705^-STAT3; STAT3, Signal Transducer and Activator of Transcription 3; pAkt, pSer^473^-Akt; Akt, protein kinase B; WT, wild-type; 1w, 1 week; 4w, 4 weeks; dox, doxycycline.

In female rNLS8 mice, WB analysis demonstrated no significant effect of either disease progression or genotype on Ob-Rb, leptin and Akt phosphorylation protein levels (Figure 4B, 4D and 5D, respectively). However, there was a significant effect of disease progression (*p*=0.028) on the phosphorylation levels of STAT3 protein (Figure 5B). Ob-Rb levels of female rNLS8 mice at 4w off-dox were significantly decreased compared to female rNLS8 mice at 1w off-dox (Figure 4B, *p*=0.016). Leptin levels of female rNLS8 mice at 4w off-dox were significantly decreased compared to age-matched WT controls (Figure 4D, *p*=0.013). STAT3 phosphorylation was significantly decreased in female rNLS8 mice at 4w off-dox compared to WT controls at 1w off-dox (Figure 5B, *p*=0.005). Akt phosphorylation levels of female rNLS8 mice at 4w off-dox were significantly increased compared to WT controls at 1w off-dox (Figure 5D, *p*=0.019).

## DISCUSSION

For many years, female subjects were largely excluded from experimental research, leading to the extrapolation of findings obtained in males to females. However, an increasing number of recent studies have demonstrated that significant sex differences exist, not only in terms of metabolism, but also in the clinical course of devastating and incurable neurodegenerative diseases such as ALS.

Although several studies have reported that ALS affects men and women differently, sex differences from a metabolic perspective remain underexplored. This is particularly notable given the growing body of evidence highlighting the role of metabolism in ALS, including the beneficial impact of higher fat reserves, which have been associated with improved survival outcomes in patients. In this study, we present the first evidence of sex-specific molecular alterations in leptin signaling pathways in the WAT, the primary site of leptin production, and the hypothalamus, a key brain region for leptin signaling, as well as in the spinal cord, a tissue profoundly affected in rNLS8 mice, a TDP-43 model of disease which is essential in dissecting mechanisms of disease as TDP-43 was identified as the major pathological protein in ∼97% of ALS [42].

In WAT, we report differences in both rNLS8 mice and WT controls, confirming sexual dimorphism. In male rNLS8 mice, there was an increase in *Ob-Rb* mRNA levels compared to WT controls, with this upregulation being more pronounced at 1w off-dox than at 4w off- dox. This increase may represent either a compensatory mechanism of adipose tissue or a disruption in the *ObR* gene splicing process. Regarding leptin mRNA levels, male rNLS8 mice exhibited a decrease at 1w off-dox, with levels further declining by 4w off-dox. This alteration in *leptin* mRNA expression levels could partly explain the changes observed in *Ob-Rb* mRNA expression, as leptin may be regulating the expression of Ob-Rb in the WAT, as it has been observed in other tissues [43]. Contrarily, in female rNLS8 mice, no differences were observed in *Ob-Rb* or *leptin* mRNA levels compared to WT controls at 1 and 4w off-dox. However, we cannot rule out the possibility that differences may be occurring later during the progression of disease in rNLS8 mice. Future experiments with longer durations following dox withdrawal are necessary to confirm this hypothesis.

On the other hand, the mRNA levels of *C/EBP*β and *PPAR*γ were increased in male rNLS8 mice compared to WT controls at 1 and 4w off-dox, along with elevated levels of *TDP-43* mRNA at 1w off-dox. These genes are primarily involved in the early stages of adipocyte differentiation, and their upregulation may reflect a compensatory attempt of this endocrine organ to restore leptin levels, which we previously observed to be decreased. In addition, both *C/EBP*β and *PPAR*γ genes are known to regulate leptin gene expression by binding to leptin promoter, directly or indirectly, during adipocyte maturation [44,45], further supporting the idea that their increased expression may be aimed at counteracting the reduced leptin signaling in the WAT observed in male rNLS8 mice. Interestingly, in female rNLS8 mice, a decrease in *C/EBP*β and *PPAR*γ mRNA expression levels was observed at 1 and 4w off-dox compared to WT controls, concomitantly with a decrease in *TDP-43* mRNA expression levels at 1w off-dox compared to WT controls. This finding may indicate that there could be alterations in WAT that affects its capacity for differentiation and expansion, even though these changes are not yet reflected in *leptin* and *Ob-Rb* mRNA expression levels, which can be also transcriptionally regulated by other factors.

In the hypothalamus of male rNLS8 mice, a decrease in *Ob-Rb* mRNA levels was observed. This is consistent with the reduced leptin expression seen in WAT, which may result in insufficient leptin reaching the hypothalamus to maintain basal levels of *Ob-Rb* expression. *POMC* mRNA levels were reduced in male rNLS8 at both 1 and 4w off-dox, with the decrease reaching statistical significance at 4w off-dox. This finding is consistent with the observed reduction in leptin levels at WAT, one of the main positive regulators of POMC neuron activity [24]. In contrast, the most remarkable findings in female rNLS8 mice were a significant increase in *AgRP* and *NPY* mRNA levels, as well as a reduction in

*POMC* levels at 4w off-dox. The reduction in *POMC* may be related to the upregulation of *AgRP* and *NPY*, since AgRP/NPY neurons release gamma-aminobutyric acid (GABA) and NPY, which inhibits POMC neurons within the arcuate nucleus of the hypothalamus [26], where all these neuronal populations are located. Although the increased expression of *AgRP* and *NPY* cannot be explained by changes in leptin levels, given that no differences in leptin expression were observed in the WAT of female rNLS8 mice, this effect may be attributed to fat mass loss, observed in mouse models of ALS, and increased energy expenditure, reported in ALS patients, which may contribute to the activation of orexigenic pathways as a compensatory mechanism to counteract the negative energy balance.

In the spinal cord, we report a downregulation of Ob-Rb and leptin protein levels in male rNLS8 mice at 4w off-dox, consistent with what we have previously described in the WAT and the hypothalamus. Additionally, a significant increase in serine phosphorylation of Akt was found in the spinal cord of male rNLS8 mice. This observation in the spinal cord is of particular interest as Akt signaling pathway plays a central role in metabolic modulating key processes such as glucose uptake, lipid metabolism, and hormonal responses to insulin and leptin [46]. Its increased could be on one hand to an increase of leptin production in other tissues, as it is also produced in the stomach, skeletal muscle and bone narrow [47,48], although in less amount than in WAT, and on the other hand to an increased in insulin levels, as it also activated this signalling pathway [49]. No differences were observed between male rNLS8 mice and WT control in tyrosine phosphorylation of STAT3.

Interestingly, in female rNLS8 mice notable findings include a decrease in Ob-Rb and leptin protein levels at 4w off-dox, along with a reduction in tyrosine phosphorylation of STAT3 at 1w off-dox. The reduction in Ob-Rb receptor protein levels aligns with the decrease in leptin levels, since Ob-Rb expression could be regulated by leptin availability [43]. The early decrease in the protein levels of the tyrosine phosphorylation of STAT3 at1w off-dox may be explained, at list in part, by the presence of leptin resistance, as it is suggested to be present in another animal model of ALS based in TDP-43 proteinopathy [50]. Further, an increase in serine phosphorylation of Akt phosphorylation levels at 1 and 4w off-dox is observed in female rNLS8 mice that could be explain as in male rNLS8 mice.

## Conclusions

In summary, our study provides evidence supporting the existence of sex differences, specifically in metabolism and alterations in leptin signaling pathway in rNLS8 mice. Future research is needed to elucidate the underlying causes of these differences, whether they are related to female sex hormones and their potential neuroprotective roles in ALS. One limitation of this study that should be taken into consideration is that in the WAT and the hypothalamus, only mRNA levels were measured, which may or may not accurately reflect protein production. Conversely, in the spinal cord, only protein levels were analysed. The presence of pathological TDP-43 aggregates observed in the spinal cord [32], could be affecting gene transcription, as TDP-43 is a protein involved in RNA processing and mRNA transport [51]. In addition, it should also be taken into consideration that this study assessed the tissues status only at 1 and 4w off-dox, so changes occurring at intermediate or later time points beyond 4w off-dox could differ from our observations.

## Declarations

### Ethics approval and consent to participate

All animal procedures were performed in accordance with the Animal Ethics Committee of the Hospital Nacional de Parapléjicos (Approval No 36OH/2019) (Spain) in accordance with the European Communities Council Directive (86/609/EEC) for the Care and Use of Animals for Scientific Purposes.

### Consent for publication

All authors have read and agreed to the published version of the manuscript.

### Availability of data and materials

The datasets used and/or analysed during the current study available from the corresponding author on reasonable request.

### Authors’ contributions

C-BC, A-FD collected the tissues; C-BC, C-FM performed statistical analysis and prepared the figures. C-FM conceived and designed the study. C-BC, C-FM drafted the manuscript. All authors critically reviewed the manuscript for intellectual content. All authors have read and agreed to the published version of the manuscript.

## Acknowledgments

The authors would like to thank Surgery Unit of the Hospital Nacional de Parapléjicos, Toledo (Spain) for their excellent technical support.

